# Aging and reproductive “rejuvenation” at a single nuclei resolution

**DOI:** 10.64898/2026.05.12.724719

**Authors:** A. C. Pearson, Jamilla Situ, Mathew Smith, Nasim Rahmatpour, Jasmine Chen, L. Y. Yampolsky

## Abstract

Aging is a multifaceted process that occurs on the background of and is significantly driven by transcriptional and cell type abundance changes. Aging-related transcriptional changes may include reduced transcription of maintenance, repair and DNA methylation genes, de-differentiation, or increased transcription of transposons. Unbiased detection of these changes and full understanding of their physiological effects requires single-cell resolution. We studied single-nuclei transcriptomes of a model microcrustacean *Daphnia magna* sampled from 3 age- and reproductive status groups: young, old reproductively senescent, and old, regaining reproductive function late in life. We detected 17 cell clusters, some identifiable as ovary or fat body-, midgut-, epithelium-, and neural tissue-related, some escaping unambiguous identification. At the same time, some well-characterized cell types were not detected, such as hemocytes or myocytes. We detect significant changes of cell type abundance with age and with reproductive “rejuvenation” in ovary- and gut-related clusters. We also detect several patterns of functional transcriptional differences between age classes with nearly all cell types, with changes between old reproductive and old non-reproductive Daphnia often reversing age-related changes, and often in good concordance with previous bulk RNAseq data.

## Introduction

Actuarial and reproductive senescence is a multifactorial process that affects many aspects of organisms’ functioning and are accompanied by profound epigenetic changes(Brunet & Rando 2017; Tarkhov et al. 2024).. A significant body of data on age-related gene expression changes largely documented in *C.elegans, Drosophila*, mice and humans. These changes include progressive dysregulation of tissue-specific transcriptional programs (“transcriptional drift” or de-differentiation) as well as increased activity of stress response and maintenance pathways that likely reflect compensatory responses to accumulating cellular damage (de Magalhães et al. 2009; Zhang et al. 2020; Gladyshev 2021; Cipriano et al. 2024; Pereira et al. 2024) However, whole body RNAseq experiments do not provide a full picture of age-related changes as both dysregulation and damage control are highly cell type-specific. Bulk transcriptomic approaches obscure this heterogeneity by averaging across diverse cell populations, thereby conflating intrinsic cellular aging with changes in tissue composition. Single-cell RNA sequencing has begun to reveal a number of patterns obscured by bulk RNAseq studies, including heterogeneity among cell lineages and age-specific remodeling (Kim et al. 2026). It has become increasingly clear that aging proceeds asynchronously across cell types, with some populations exhibiting pronounced functional decline while others remain comparatively stable or even undergo compensatory remodeling (Angelidis et al. 2019; Tabula Muris Consortium 2020). Single-nucleus RNA sequencing extends these capabilities to tissues and physiological states that are difficult to access using whole-cell approaches, while providing a direct view of transcriptional regulation through measurement of nascent nuclear transcripts. These features make single-cell and single-nucleus transcriptomics uniquely suited to addressing one of the central unresolved questions in aging biology: how aging-related decline can be partially reversed during reproduction. In diverse organisms, reproduction is associated with restoration of somatic function and extended survival, implying the existence of endogenous rejuvenation mechanisms. Identifying these mechanisms requires resolving how transcriptional states change within specific cell types during aging and reproductive transitions. By enabling direct comparison of cellular transcriptional states across physiological conditions, single-cell and single-nucleus approaches provide a powerful framework for uncovering the cellular basis of aging and reproductive rejuvenation.

Aging has traditionally been viewed as a unidirectional process of progressive cellular decline; however, accumulating evidence suggests that differentiated cells can undergo partial reprogramming under physiological conditions, altering gene expression profiles in ways that restore functional capacity or regenerative potential. Such transitions have been observed in diverse systems, including regeneration, stem cell activation, and reproductive cycles, and are often accompanied by coordinated changes in chromatin organization and transcriptional regulation (Rando & Chang 2012; Lu et al. 2020). Single-cell and single-nucleus transcriptomic approaches provide a powerful means to detect such transitions, as they allow identification of intermediate or hybrid transcriptional states that would be obscured in bulk measurements. In organisms such as Daphnia, where reproduction is associated with restoration of somatic performance, de-differentiation or partial reprogramming of specific cell populations may represent a key mechanism linking reproductive state to systemic rejuvenation. Resolving whether such transcriptional plasticity occurs during aging and reproduction requires direct characterization of cell-type-specific transcriptional states, which single-nucleus transcriptomics uniquely enables.

Another mechanism increasingly implicated in aging and rejuvenation involves activation of transposable elements (TEs), which are normally repressed by epigenetic mechanisms but can become transcriptionally active during aging. TE activation has been documented across diverse taxa, including flies, mammals, and other eukaryotes, and is thought to reflect age-associated deterioration of chromatin maintenance and genome surveillance systems (Li et al. 2013; De Cecco et al. 2013; Wood et al. 2016; Della Valle et al. 2025; Merenciano et al. 2025; Lemus et al. 2026). Activation of TEs can contribute to genomic instability, alter gene regulation, and trigger innate immune responses, thereby promoting cellular dysfunction. This TE activation can be seen as both a consequence and a cause of aging (Merenciano et al. 2025). Because TE activation and repression can vary substantially across cell types, bulk transcriptomic approaches provide only limited insight into their role in aging. Single-cell and single-nucleus RNA sequencing offer the opportunity to identify specific cell populations in which TE expression changes during aging or reproductive transitions, providing a more precise understanding of how genome regulatory stability contributes to aging and rejuvenation. Such analyses may help clarify whether reproductive rejuvenation involves restoration of epigenetic control over transposable elements and stabilization of cellular transcriptional programs.*Daphnia* as a model organism for longevity studies has a number of advantages. It’s mode of reproduction, cyclic parthenogenesis, allows maintaining genetically uniform, and yet outbred clones, thus controlling for heterogeneity within cohorts without the detrimental effects of inbreeding and providing a possibility to test the same genotype in different environments. Daphnia’s transparent body allows in-vivo determining ovary cycle and embryonic development stages and fecundity, making it easier to evaluate age-specific reproductive status. Finally, rapidly developing genomics resources provide tools for referencing and annotating genomic data. *Daphnia’s* phylogenetic position close to the root of Pancrustacea bridges the gap between better studied ecdysozoan models such as *Caenorhabditis* and *Drosophila.* On the other hand, the disadvantage of *Daphnia* as a model for single cell transcriptomics is that, currently, there is no consistent methodology for in-situ hybridization for adult *Daphnia*, despite several promising attempts (Sagawa et al. 2005;Liu et al. 2014), making tissue annotation a challenge.

To expand our understanding beyond the handful of model systems and to generalize DE patterns across multiple organisms we report single-nuclei RNAseq experiment comparing young *Daphnia* to old, reproductive senescent ones and, additionally, old, reproductive senescent *Daphnia* to similarly old, but reproductively “rejuvenated” (Dua et al. 2025) ones. Reproductively “rejuvenated” *Daphnia* show post-senescence rebound of asexual reproduction (often with clutch sized exceeding those produced by younf females) and demonstrate a distinct transcriptomics state, with many genes showing reversal of age-related expression changes. These include paralogs of apolipoprotein D (homologs of *Drosophila nlaz),* liposomal lipases, peptidases, peroxidases and cytochromes, among others (Dua et al 2025). Here we report similar differential expression comparisons at a single-nuclei resolution.

## Materials and Methods

### Daphnia provenance and maintenance

The experiment was conducted using a laboratory parthenogenetic clone of *D. magna* GB-EL75-69 (Basel University *Daphnia* Stock Collection, Basel, Switzerland) originated from a permanent lake in London, UK and maintained in the lab for over 10 years. The stock of this clone was in ADaM reconstructed pond water medium (Klüttgen et al. 1994) in a 20 °C, 12:12 light:dark photoperiod incubator, fed with green alga *Tetradesmus obliquus* (Turpin) M.J.Wynne culture (synonym: *Scenedesmus acutus*).

Grandmothers of experimental animals were sampled from the stocks and maintained in groups of 5 in 100 mL jars equipped with 50 mL, 1 mm^2^ bottom mesh inserts that allow easy removal of neonates at each water change (Beam et al. 2024). These founding individuals and their daughters were maintained at the density of 5 individuals per 100 mL of ADaM medium with *Scenedesmus acutus* culture added as food added daily in the amount of 1E6 cells per mL and water replaced and newborn removed every 4 days, at the same temperature and photoperiod as described above.

Four separate cohorts staggered 10-25 days apart were established from the second (mother) generation maintained at the conditions described above by sampling female neonates age <24 hours born to mothers 15-50 days old. These juveniles were kept in groups of 10 in jars containing 100 mL of ADaM medium, fed in the same manner, until day 6, at which point they were separated into individual vials containing 20 mL of ADaM medium. Food was added to these vials daily in the amount of 1E6 cells/mL and water was replaced at every new brood of neonates produced, or at each molting event in case no neonates were produced in a given ovary/molting cycle (i.e., every 3-4 days). Number of neonates produced in each clutch was recorded and mortality of cohort individuals was recorded every day.

At the age 15-20 days the “young” (“Yox” thereafter) samples were collected, one sample per cohort, each containing 15 individuals. Additional back-up samples were also collected. As the cohort individuals reached the age of 90 days the “old, non-reproducing” and “old, reproducing samples” were collected, (“ONx” and “ORx” thereafter, respectively). Criteria of “reproductive rejuvenation” by which individuals were added to the ORx samples were similar to those in Dua et al. 2024: at least 2 skipped clutches prior to production of a clutch of at least 6 individuals. Individuals with at least one skipped clutch and current clutch of less than 3 eggs (typically 0 or 1) were collected into the ONx samples. In all three types of samples individuals within 24 hours of molting were sampled to assure uniformity of the ovary/molting cycle phase and any eggs present in the brood chamber were removed to assure that only somatic tissues are sampled and to reduce ambient RNA amounts. A mock egg removal procedure was performed on individuals carrying no eggs in the brood chamber. Females were frozen in liquid nitrogen immediately after that and stored at -80 °C until single nuclei library prep.

### Single nuclei isolation and sequencing library preparation

Steps described in this section were performed at Abiosciences, Inc lab according to the procedure optimized for *Daphnia* single nuclei RNAseq library preparation (Abiosciences 2025). Briefly, 15-20 frozen females were combined into each library replicate, single nuclei were isolated using NIM*+FACS procedure (McLaughlin et al. 2022) with frozen *Daphnia* homogenized in a Dounce homogenizer in nuclei isolation buffer with Triton-X, followed by FACSorting (Sony MA900) using PI and DAPI as nuclei markers. Sorted nuclei were stained with Acridine Orange/PI and counted using Revvity Cellaca MX cell counting equipment. Libraries were prepared using 10X Genomics Chromium Single Cell 5’ v2 protocol, cleaned up using double-sided SPRI cleanup protocol and sequenced on an Illumina NovaSeq X Plus platform.

### Bioinformatics

Raw reads were processed by CellRanger v8 (including intronic reads; Zheng et al. 2017) using *D. magna* genome assembly GCA_040143795.1 (D. Ebert and P.Fields, personal communication). The resulting filtered UMI abundance matrices were combined into a single dataset, normalized, doublet-removed and analyzed for cluster composition, cluster markers and within cluster differential expression using Seurat Ver. 5.3.0 (Hao et al 2023) using UMAP as the dimensionality reduction algorithm and the default clustering algorithm. Seurat scripts are available in the Supplementary materials. Briefly, the following parameters were used other than defaults: cells filtering: **min.features** = 200; feature filtering: **min_cells** = 50; doublet removal expected rate = 0.05, clustering: npcs = 50, **dims** = 1:30, **k.param** = 20, **resolution** = 0.6; markers identification and differential expression: **test.use** = “MAST”, **logfc.threshold** = 0.25, **min.pct** = 0.1. See Supplementary Tables S1 and S2 for doublet removal results and the effects of clustering parameters on the number of clusters.

Cluster annotation was accomplished by matching markers to previously published *D.magna* single-cell transcriptomics dataset (Krishnan et al. 2025) and to *Drosophila* Fly Cell Atlas using two complementary enrichment approaches. First, we tested for the enrichment of shared markers relative to background expectation using Fisher Exact Test, with the Universe of genes eligible to become markers set by the same parameters used for marker identification. Second, we propose a simple rank-aware enrichment test that aims to detect non-randomness in ranks of markers between the annotated and reference sets, specifically relying on genes that have unusually high rank in both lists of markers (Appendix 1).

Gene Ontology (GO) enrichment analysis for clusters’ markers and for genes differentially expressed within clusters was conducted using Fisher Exact Test with the Universe determined as the number of genes in the reference dataset eligible to be detected as markers or DE genes by surviving Seurat filtering. For cluster’s markers this was defined as the smaller (more conservative) of the genes surviving the threshold of **min.pct** = 0.1 in the smallest of clusters and genes that were detected in at least 20 cells in SCT-transformed dataset., yielding the universe of 3925 genes. For up- or downregulated differentially expressed genes within clusters this was further reduced to only include genes with GO annotation, resulting in the universe of 1028. This is conservative, because finding a match between DE genes’s GO terms and reference GO terms is conditional on GO annotation availability.

To assess whether cells derived from Yo, On, or Or treatments were over-represented in each cluster, we analyzed the abundance of cells from each age class within each cluster using a sample-level negative binomial generalized linear model (NB-GLM) with a log-offset for total cell count. This approach tests for treatment-of-origin differences within clusters while accounting for sample-to-sample variability (batch effects) among biological replicates within treatments. P-values were corrected for multiple testing using the Benjamini–Hochberg false discovery rate (FDR) procedure.

### AI use

ChatGPT v.4 was consulted at the stage of bioinformatic analysis, including writing and debugging R scripts and ChatGPT v.5 was used at the stage of manuscript editing.

## Results

### Doublet removal, clustering, and cluster annotation

Results of doublet removal procedure are presented in Supplementary Table S1. Between 7% and 10% of cells were deemed doublets and removed from the analysis; the rate did differ among samples (Log Likel-hood c2 = 19.9, P<0.006), as well as between Young and Old samples (8.48% vs 9.76%, respectively, 2-tailed Fisher’s Exact Test P<0.012), but no differences between reproducing (Or) and non-reproducing (On) samples within the Old age class (9.45% vs. 9.97% respectively, 2-tailed Fisher’s Exact Test P>0.63) and no correlation between the number of cells in the raw data and percentage of doublets (linear regression -3.06E-6, P>0.59). This is consistent with modest but statistically significant biological age–specific differences in aggregation propensity, or nuclear integrity that influence encapsulation efficiency. However, the small magnitude of this effect, together with the lack of association with library size or reproductive status, indicates that the majority of doublet occurrences are driven by stochastic co-encapsulation events during nuclei partitioning rather than by systematic biological or technical factors.

Cycling over clustering parameters revealed no additional benefit in increasing resolution or the number of PCA axes to be utilized beyond the values of **res**=0.4 and dims=1:25 (Supplementary Table S2) in terms of our ability to annotate cell clusters, due to oversplitting otherwise well-defined clusters.

Summaries of data on clusters obtained with resolution=0.4 are presented in Tables 1 and 2. Median depth per cluster varied between 404 and 1051, with overall median depths being 580 UMIs. Table 1 presents the summary of annotations by matching clusters to previously described *Daphnia* cell types annotated in our resent single-cell transcriptomics study. Marker list enrichment analysis demonstrated a radical difference in single nuclei vs. single cell clustering, resulting in difficulty annotating cluster unambiguously. In particular the annotation of two cell-rich clusters 0 and 5 (Fig. 1A is ambiguous with respect matching them to either ovary or fat body: cluster 0 is matched to ovary and ovary-associated hemocytes in the *DaphniaSN/DaphniaSC* match and to fat body in a *Daphnia*SN / *Drosophila* match, while cluster 5 was matched to fat body in both enrichment analyses. Several clusters identified in this study significantly matched to cell types described in Krishnan et al. (2026) that escaped a non-ambiguous annotation in our previous study. For example, the well-defined cluster 1 (Fig. 1, Table 1 is significantly matched to Krishnan et al cluster 16, which, in turn, has affinities to such diverse *Drosophila* cell types gut, fat body, neurons, and myocyte, but failed to be significantly matched to any *Drosophila* cell type in this study.

**Fig. 1.**
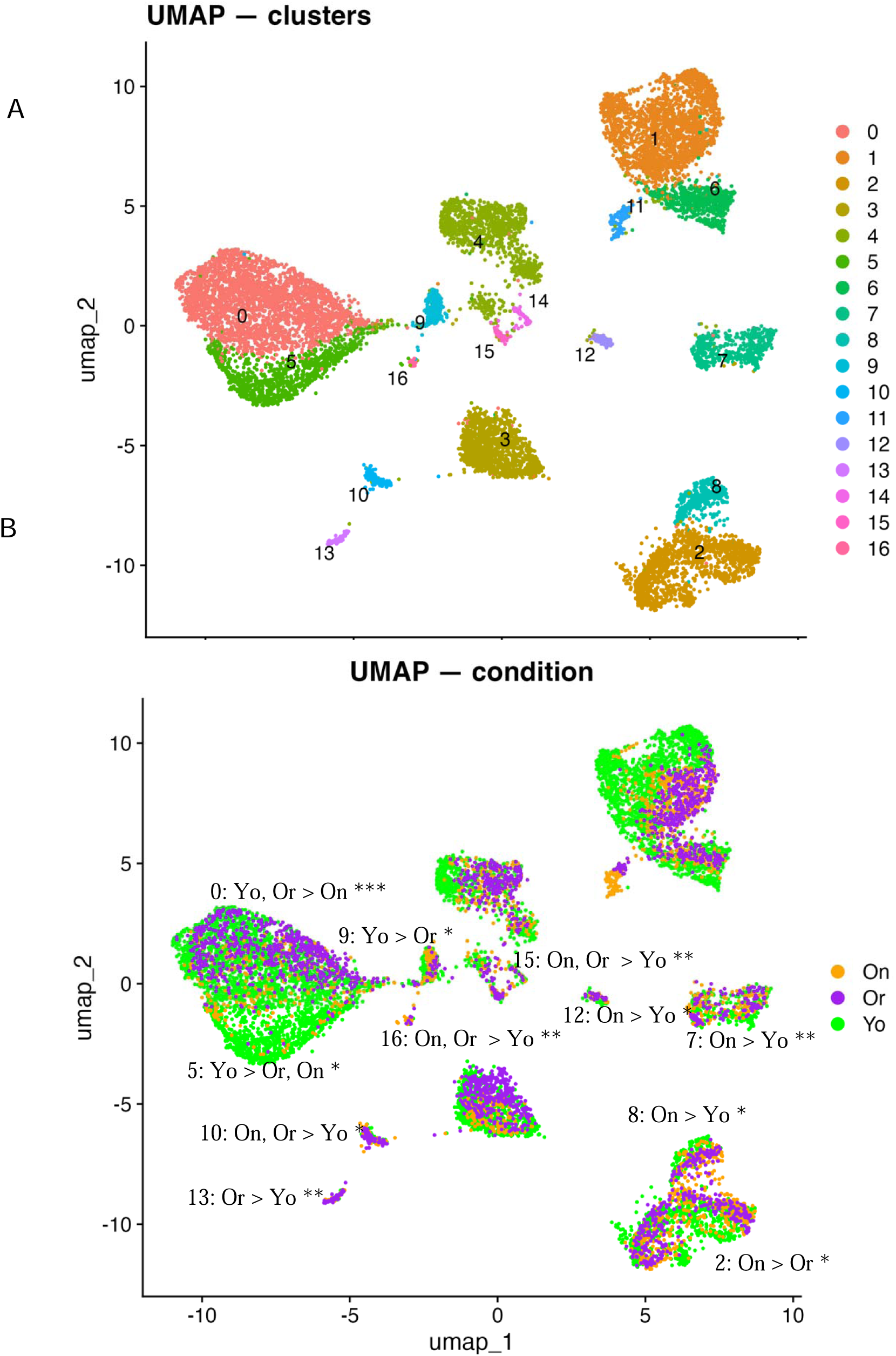
A: cells colored by cluster; B: cells colored by age / reproductive status. Here and throughout Green: young, orange: old, non-reproducing, purple: old, reproducing *Daphnia*

**Table 1.** Summary of annotation by rank-product enrichment test. Matches with the lowest FRD values shown; matches for artefactual clusters reported in Krishnan et al. 2026 omitted. See Supplementary data for compete list of matches.

| cluster | Inferred annotation | Matched to markers of Krishnan et al. (2026) clusters |  | Matched to FCA group markers |  |  |
| --- | --- | --- | --- | --- | --- | --- |
|  |  | ID | Krishnan et al. annotation | Min Enrichment FDR | tissues or organs | Min Enrichment FDR |
| 0 | Ovary | 19 | ovary | 7.62E-16 | fat body | 4.96E-03 |
|  |  | 26 | ovary-associated hemocytes | 7.41E-11 |  |  |
| 1 | Ambiguous | 16 | ambiguous: gut / fat body / neural / myocyte | 1.68E-177 | -- | -- |
| 2 | Ambiguous (possibly: neural tissue) | 17 | ambiguous: neural / crop / fat body | 2.80E-04 | glia<br>neural tissue<br>follicle<br>pylorus/hindgut | 4.25E-43<br>7.30E-04<br>1.34E-02<br>1.57E-03 |
| 3 | Midgut | 12 | Midgut | 9.65E-06 | midgut | 7.59E-04 |
| 4 | Epithelium (possibly fore- or hindgut) | 27<br>21 | Foregut or hindgut<br>Epithelium | 5.76E-05<br>6.03E-02 | -- | -- |
| 5 | Fat body / non circulating hemocytes | 5 | Fat body-like non-circulating hemocytes | 4.76E-11 | fat body | 0.1 |
| 6 | Ambiguous (possibly: neural tissue) | 16 | ambiguous: gut / fat body / neural / myocyte | 5.58E-03 | neural tissue<br>ovary<br>hemocyte | 8.71E-47<br>5.61E-12<br>4.20E-05 |
| 7 | None | -- | -- | -- | -- | -- |
| 8 | Neural tissue | 17 | ambiguous: neural / crop / fat body | 1.01E-05 | neural tissue<br>glia | 4.11E-97<br>2.50E-40 |
| 9 | Ambiguous (possibly: | 14 | ambiguous: follicle / myocyte / | 3.65E-34 | neural tissue<br>ovary<br>trachea | 1.81E-09<br>4.52E-07<br>0.01 |
|  | neural<br>tissue) |  | neural /<br>Malpighian<br>tubules |  |  |  |
| 10 | Midgut | -- | -- | -- | midgut | 1.46E-04 |
|  | Table 1,<br>continued |  |  |  |  |  |
| 11 | None | -- | -- | -- | -- | -- |
| 12 | Neural<br>tissue | -- | -- | -- | neural tissue | 1.94E-05 |
| 13 | Ambiguou<br>s | 33 | ambiguous:<br>esophagus / fat<br>body / germ line<br>/ non-neural<br>sensory /<br>midgut | 2.76E-45 | -- | -- |
| 14 | Follicle | 24 | ambiguous:<br>follicle / fat body<br>/ neural | 3.33E-05 | Follicle | 5.72E-03 |
| 15 | Neural<br>tissue | -- | -- | -- | neural tissue | 2.55E-03 |
| 16 | Ambiguou<br>s<br>(possibly:<br>ovary) | 34 | Epithelium | 0.075 | ovary | 3.10E-50 |

**Table 2.** Frequencies of genes with co-directional (both down or both up) and compensatory/reversal DE in old, nonreproducing vs. young (aging) and old, reproducing vs. old, nonreproducing (“rejuvenation”).

| cluster | DE both | Down in both | Up in both | Down in aging<br>Up in<br>“rejuvenation” | Up in aging<br>Down in<br>“rejuvenation” | Bonferroni-<br>adjusted FET |
| --- | --- | --- | --- | --- | --- | --- |
| 0 | 81 | 2 | 0 | 39 | 40 | 3.28e-20 |
| 1 | 20 | 2 | 4 | 8 | 6 | 0.67 |
| 2 | 20 | 4 | 3 | 9 | 4 | 1 |
| 3 | 58 | 0 | 1 | 23 | 34 | 2.16e-14 |
| 4 | 8 | 3 | 1 | 2 | 2 | 1 |
| 6 | 3 | 1 | 0 | 0 | 2 | n/a |
| 7 | 41 | 4 | 0 | 23 | 14 | 6.94e-07 |
| 8 | 6 | 0 | 0 | 4 | 2 | 0.53 |

Despite functional annotation difficulties, clusters are well-defined in terms of specific markers (Fig. 2). Clusters 0 and 5 with overlapping functional annotation as ovary / fat body related share markers with each other (TEP2/Macroglobulin, SANS/Ankyrin repeat containing protein, CAR/ Superoxide dismutase, collagen) and with a small cluster 16 (YL/ Low density lipoprotein receptor, an ovary marker in *Drosophila).* The same collagen marker is also shared between both clusters 0 and 5 and cluster 14 annotated a follicle cells. Some of clusters provisionally identified as various neuron types also share markers, for examples cluster 15 and 6 share PDE1C / 3’,5’-cyclic-GMP phosphodiesterase, which also has high expression in *Drosophila* neurons. Likewise, clusters 2 and 9 share paralogs of *Drosophila* GAD1, a nervous system-specific glutamic acid decarboxylase, while clusters 8 and 15 shared a marker homologous to *Drosophila SMYDA-5* protein with high expression in neurons, myocytes and epithelium. Some clusters that do not appear to be functionally related also share markers that are shared markers between *Drosophila* cell types: clusters 14 (follicle cells) and 9 (neurons) share an ortholog of *Drosophila UNC* with known expression in reproductive neural system.

**Fig. 2.**
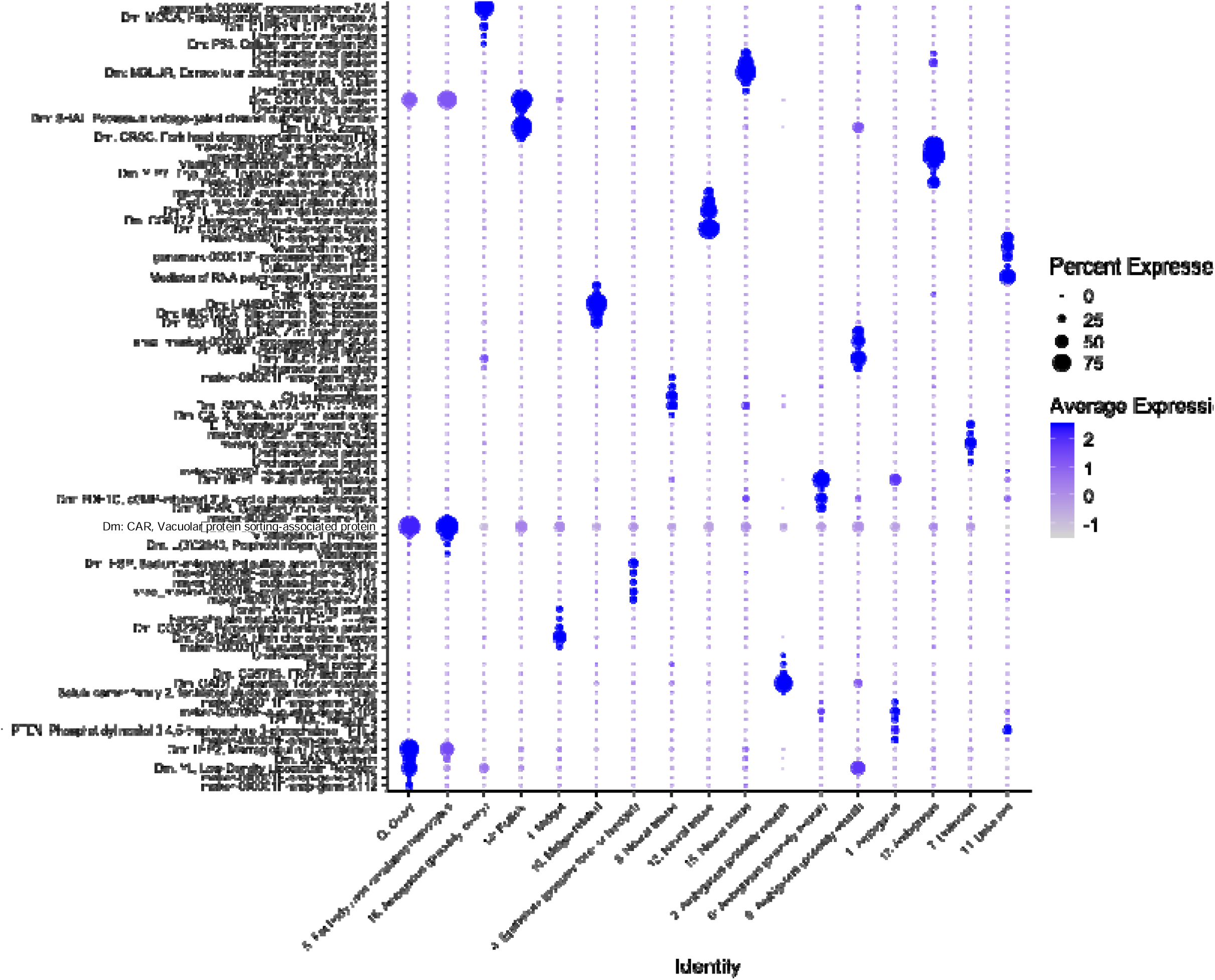
Dotplot of 5 top markers of 17 clusters. Horizontal axis: clusters annotated as on Table 1 and sorted by functional properties of inferred annotations. Vertical axis: markers labeled by *Drosophila* ortholog (Dm:; where available) and brief functional annotation; when neither is available the DM3 gene ID is used. See supplementary materials for the full list of markers.

### Cluster composition

Several clusters significantly differed in cell composition with respect to the three age and reproductive status classes in replicate-aware test (Fig. 1B, Fig.3, and Supplementary Table S3). Not surprisingly, clusters 0 and 5 (presumed ovary cells and fat body, but not cluster 14 identified as follicle cells) contained disproportionally large number of cells from young samples, relative to both old reproducing and old non-reproducing ones, as did cluster 9 provisionally identified as a type of neurons. Sevel other clusters were enriched in cells from old age samples, including the unidentified clusters 2, 7, and 16 and different neural tissue clusters 8, 12, and 15. Of the two midgut-related clusters, the larger cluster 3 showed enrichment of cells from old, reproductively active samples relative to old reproductively senescent ones, while the other, smaller cluster 10 contained disproportionally high cell count from old samples independent form their reproductive status.

**Fig. 3.**
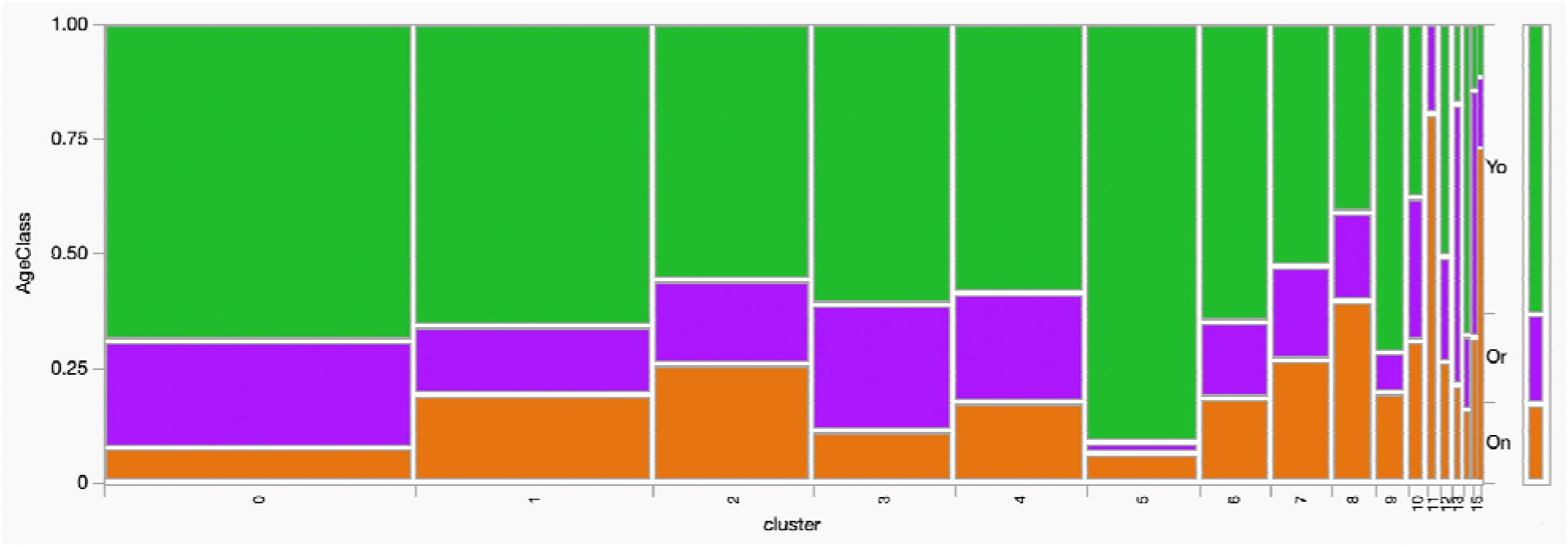
Cluster composition by age class. Bar on the far right represents whole-library counts. Log-likelihood ration c2=1803.1, P<0.0001. All clusters except 7 and 17 differ in Age class composition. See Supplementary Table S3 for full statistics.

### Differential expression within clusters: Old, non-reproducing vs. young Daphnia

Thirteen out of 17 clusters showed at least some genes with aging-related differential expression (Fig. 1B, Supplementary Data). None of the GO terms showed any significant enrichment, except for GMP synthesis functionality (see below), but revigo plot reveals some association between GO semantics and cluster identity (Supplementary Fig. S1). Many individual genes showed either up- or down-regulation in the “old” samples in individual clusters. Cluster 0 (presumed ovary cells) showed downregulation of several structural proteins (collagen, cuticular protein), several proteins with mitochondrial localization (ferritin, malate dehydrogenase), an ortholog of *Drosophila **14-3-3e*** involved in Ras/MAPK cascade including regulation of germ cell migration, gonad formation, a cathepsin whose human and *Drosophila* orthologs may be involved in protein degradation and apoptosis, and at least one enzyme in glutathione synthesis pathway (cystathionine gamma-lyase). The same cluster showed aging-related upregulation of three different homeobox proteins, a mitochondrial cytochrome P450 homolog, and one of several genes identified to be of transposon or retrovirus origin (polyprotein encoding). Functionally related cluster 5 (presumed fat body) showed downregulation of the same ortholog of ***14-3-3e***, an ortholog of *Drosophila **Baldspot*** very long chain fatty acids elongase, an ortholog of *Drosophila* ovary marker ***Mael*** involved in transposon suppression during oogenesis, a glycogen synthase and one of several vitellogenin precursors, among others. The same cluster showed up-regulation in “old” samples of a different polyprotein-encoding gene of presumed retroviral origin, several mitochondrial enzymes, a partially overlapping set of homeobox proteins with those up-regulated in cluster 0 and, unexpectedly, the same ferritin homolog that showed down-regulation in cluster 0.

Predictably, clusters identified as midgut-related (3 and 10) both lost expression of a variety of proteinases and gained expression of several membrane transported proteins, solute carriers and membrane signaling proteins (Aquaporin and Semaphorin homologs, among others) and one of many paralogs of GMP synthases, orthologous to *Drosophila **bur*** (see below). Cluster 4 (presumed epithelium cells, possibly fore- or hindgut) showed downregulation of a superoxide dismutase and two different Zn finger proteins and up-regulation of a cystathionine gamma-lyase paralog and of a different paralog of gmp synthase.

We could not detect any consistent patterns in up- or-down-regulation of transcripts abundance in presumed neural tissues. Cluster 8 showed down-regulation of a cGMP-specific 3’,5’-cyclic phosphodiesterase involved, in *Drosophila* in regulation of intracellular concentration of cGMP and, consistent with other clusters, up-regulation of a transposon/retroviral polyprotein encoding gene.

Among clusters for which we were unable to obtain an unambiguous functional annotation, cluster 1 stood out as showing aging-related differential expression consistent with other clusters and previous whole-body data. It lost, in “old” samples, expression of the same cGMP-specific phosphodiesterase that was also down-regulated in cluster 8, the homolog of ***14-3-3e*** also down-regulated in clusters 0 and 5, superoxide dismutase down-regulated in clusters 5 and 9 (among others), and cystathionine gamma-lyase down-regulated in clusters 0, 3, and 6. The “old” samples within this cluster gained expression of different cytochrome P450, several genes of transposon/retroviral origin, and, notably, three different paralogs of GMP synthase, two of which are the same that are up-regulated in clusters 4 and 6.

### Reversal of aging-related differential expression in old, reproducing Daphnia

Eight clusters showed reversal of gene expression associated with “rejuvenated” condition (Table 2, Supplementary Table S4). Reversal of gene expression was defined as identifying cases in which the expression of a gene was statistically similar in “young” (“Y”) and “old, reproducing” (“OR”) samples, but statistically different in “old, nonreproducing” (“ON”) samples, indicating that the ON samples were following a “younger” expression pattern. In those clusters in which any genes showed a significant “rejuvenation” DE (OR vs. ON), the genes with aging DE were overwhelmingly overrepresented among genes with “rejuvenation” DE (Table 3) indicating that transcriptional changes in reproductively “rejuvenated” *Daphnia* were dominated by reversing or preventing aging-related changes. A striking exception was cluster 5 (presumed fat body-related), in which not a single gene showed “rejuvenation” DE, in contrast to functionally related cluster 0 with 132 such genes, due to very low cell counts in On and Or samples in this cluster, preventing meaningful DE analysis. Furthermore, among genes with significant DE in both comparisons, reversals of age-related changes dominated over co-directional changes (Table 2). Of course this analysis is inherently biased via regression to the mean bias, as genes with unusually high expression in “old, non-reproducing” samples are predestined to have lower, not higher expression in the “old, reproducing” samples. Therefore, this analysis should be interpreted solely as an indication of old, reproducing samples being different from old, non-reproducing ones and for a comparison of frequencies of genes that reverse or prevent age-related up-regulation to those that reverse or prevent age-related down-regulation. In particular, clusters 0 (presumed ovaries), 3 (presumed midgut) and 7 (unknown) showed higher than expected counts of genes with either reversal of aging-related DE by down- or by up-regulation in the “old, reproducing” samples, in roughly equal frequencies each (Table 2).

**Table 3.** Enrichment of genes with “rejuvenation” DE among genes with aging DE (out of total of 1505 eligible genes). Data for all DE shown regardless of directionality. “n/a” – not applicable due to no genes with “rejuvenation” DE.

| cluster | DE<br>On.vs.Yo | DE<br>On.vs.Or | DE both | Bonferroni-<br>adjusted FET |
| --- | --- | --- | --- | --- |
| 0 | 258 | 132 | 81 | 1.43e-32 |
| 1 | 94 | 52 | 20 | 4.76e-11 |
| 2 | 91 | 59 | 20 | 3.91e-10 |
| 3 | 110 | 112 | 58 | 2.55e-41 |
| 4 | 21 | 31 | 8 | 3.55e-08 |
| 5 | 271 | 0 | 0 | n/a |
| 6 | 30 | 4 | 3 | 4.81e-4 |
| 7 | 142 | 71 | 41 | 5.10e-25 |
| 8 | 36 | 9 | 6 | 1.65e-07 |
| 9 | 4 | 0 | 0 | n/a |
| 10 | 19 | 0 | 0 | n/a |
| 11 | 0 | 87 | 0 | n/a |
| 12 | 10 | 0 | 0 | n/a |
| 13 | 0 | 3 | 0 | n/a |
| 15 | 0 | 2 | 0 | n/a |

Within cluster 0, the structural proteins (collagen, cuticular proteins, among others) which had been seen to be downregulated in ON samples were upregulated in OR samples. Similarly, mitochondrial proteins such as ferritin also underwent reversal of age-related down-regulation in OR, as well as glutathione biosynthesis enzymes such as cystathionine gamma-lyase. The three homeobox proteins seen to be upregulated in ON were downregulated in OR, as was a cytochrome P450, and an ortholog of *Drosophila **sicily*** (known to be involved in the function of electron transport components). Some, but not all, of the proteinases in cluster 3 (presumed midgut) seen to lose activity in ON samples displayed upregulation in OR.

In cluster 1 (lacking an unambiguous functional annotation), the cGMP-specific phosphodiesterase which was seen to be downregulated in ON samples was upregulated in OR samples, although interestingly, it did not undergo the same reversal in cluster 8, where it was also lost in ON. The same cystathionine gamma-lyase undergoing reversal in cluster 0 also experienced reversal in cluster 1, but not in clusters 3 or 6. Similarly, ambiguous cluster 2 did not contain any detectable expression patterns.

While reversals of DE in old, non-reproducing samples are interesting, significant co-directions changes in old, reproducing samples may be of special interest as well. If DE changes in senescent *Daphnia* reflect protective responses against age-related changes, extending these responses in old-reproducing *Daphnia* (further up from already up-regulated genes, or, perhaps less likely, further down from an already down-regulated level) may be a mechanism of retaining (or re-gaining) reproductive function. Such co-directional changes are less common (Table 2, Fig. 4, Supplementary Table S5) and there is a significant heterogeneity among clusters in terms of how common they are: likelihood ratio c2= 60.4 P<1.12e-5, dominated by higher than by chance gene count for “down in both” in cluster 4 and ‘up in both in cluster 1.

**Fig. 4.**
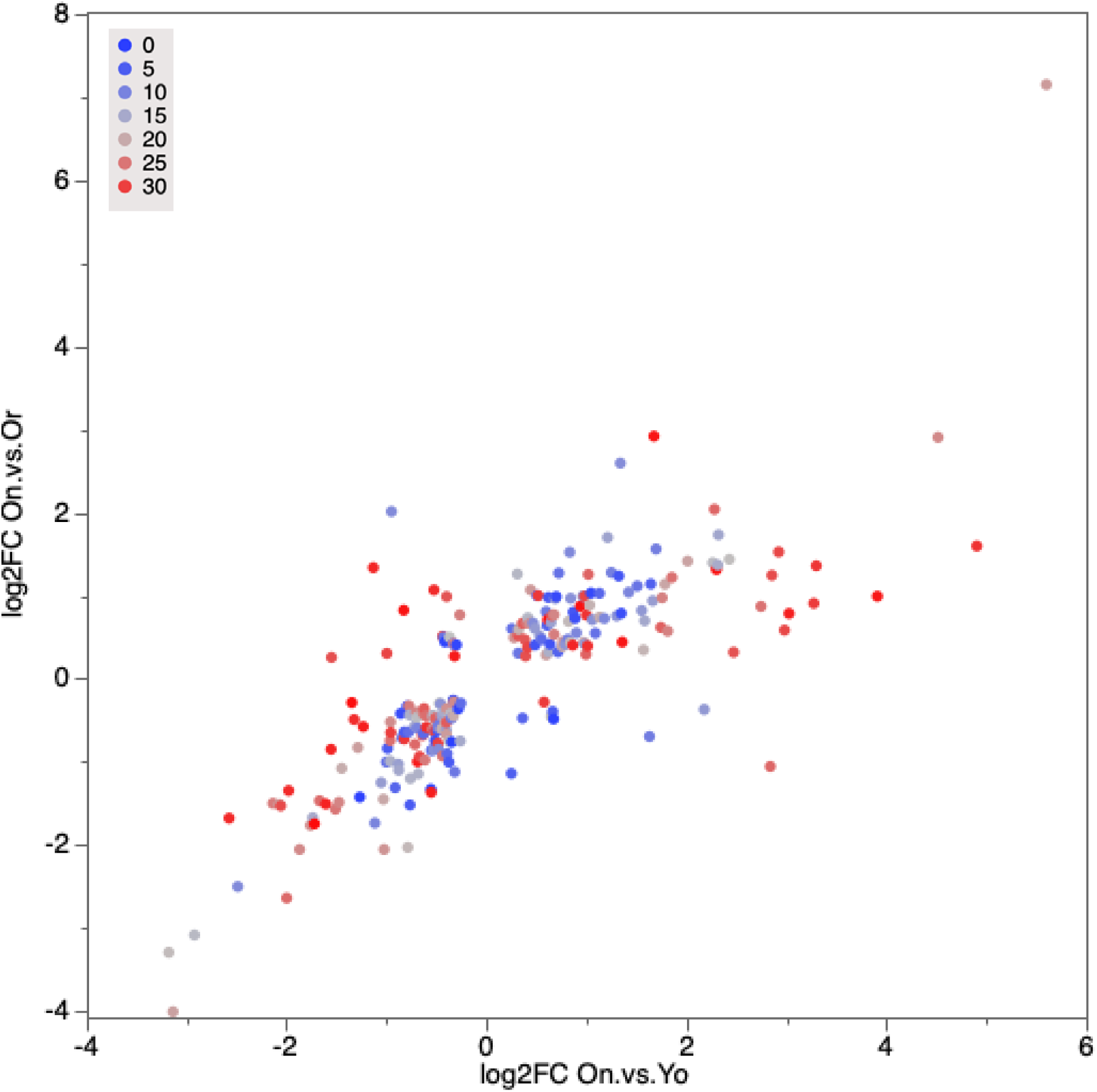
A 2-D volcano plot showing DE (log_2_(fold change)) in the comparison of old, nonreproducing (On) *Daphnia* to young (Yo) *Daphnia* (horizontal axis) and of old, nonreproducing *Daphnia* to old, reproducing (Or) *Daphnia* (vertical axis). Each data point is a gene, painted by -log_10_(P_adj_) in the On vs. Yo comparison. Only genes that pass the filters (see Methods) and show a significant (P_adj_<0.1) DE in both comparisons are shown.

Several notable proteins followed the “both Up” or “both Down” patterns. In unidentified cluster 1 the reproductively “rejuvenated” *Daphnia* further up-regulated already up-regulated in non-reproducing old *Daphnia* the homolog or *Drosophila **CYP18A1**, a* P450 enzyme that inactivates 20-hydroxyecdysone hormone which is known to suppress molting and ovulation (Sumia et al. 2014; see discussion). Similar pattern in the same cluster was observed in the homologs of *Drosophila **DIP-beta*** and ***Ca-***α***1D***, 2 membrane proteins with expression in the nervous system, along with one protein of unknown function.

The same pattern in cluster 2 was observed in two proteins both implicated in actin cytoskeleton reorganization, phosphatase-and-actin regulator and StAR-related lipid transfer 13 protein, and in a homolog of human Zinc finger C2HC domain-containing protein 1A, whose *Drosophila* ortholog is known to be highly expressed in male reproductive system.

Genes showing “both Down” DE in the two comparisons were more numerous (Supplementary Table S5). Some of the inferred function of genes in this categorically appeared paradoxical. For example, “both Down” pattern in cluster 0 (presumed ovary) was observed in the homolog of *Drosophila **mael,*** a gene with a well-known function in oogenesis, namely in repressing transposable elements in the germline. While not surprising that this function declined in reproductively senescent *Daphnia,* the fact that it further declined in reproductively “rejuvenated” ones is counter-intuitive (see Discussion). Other notable cases were more predictable. For example, the homolog of *Drosophila **crol*** was down in old, non-reproducing *Daphnia* and further down in “rejuvenated” ones in cluster 4 (presumed epithelium). In *Drosophila* the transcription factor ***crol*** has non-exclusively high expression in epithelial cells, is involved in regulation of cell cycle and is known to be induced by 20-hydroxyecdysone. Thus its widespread down-regulation is consistent with the “both Up pattern” of 20-hydroxyecdysone inactivator ***CYP18A1*** in a different cell type. Another example of “both Down” DE pattern consistent with enhanced anti-aging activity in the “rejuvenated” *Daphnia* was the observed in two different copies of retroviral polyprotein in the unidentified cluster 7 (see Discussion).

## Discussion

### Cluster annotation difficulties

Single cell (SC) and single-nuclei (CN) transcriptomics in *Daphnia* is in its infancy. We have recently published (Krishnan et al. 2025) a survey or SC transcriptomics in *D.magna* identifying several key cell types in adult *Daphnia,* including neurons, myocytes, fat body cells, epithelium cells and female- and male-specific reproductive system cells. Yet, we failed to functionally annotate many well-defined clusters due to their markers being highly *Daphnia-*specific or, even when reliable *Drosophila* or human orthology could be established, due to ambiguous matches to annotated cell types in these organisms. The greatest obstacle to interpreting the results of this study is similar. Indeed, with the exception of presumed ovaries, fat body / fat body–associated hemocytes, epithelium, and midgut, most clusters escaped unambiguous identification. Some clusters showed no shared markers with the scRNA-seq clusters of Krishnan et al. (2025), while others, such as cluster 1 in the present study, shared markers with a well-defined Krishnan et al. cluster (cluster 16) that itself could not be conclusively annotated. Still other clusters showed significant marker overlap with Krishnan et al. clusters that were deemed artefactual in that analysis. Similarly, few clusters showed strong marker correspondence with Drosophila Fly Cell Atlas scRNA-seq cell types. In principle, the lack of concordance between snRNA-seq and scRNA-seq cell clustering is not unexpected. Both biological and technical factors contribute to such discrepancies. Biologically, snRNA-seq primarily measures nuclear RNA, including nascent and incompletely spliced transcripts, and therefore reflects ongoing transcriptional activity, whereas scRNA-seq captures both nuclear and cytoplasmic RNA and thus reflects the accumulated steady-state transcript pool (Lake et al., 2016; Bakken et al., 2018; Slyper et al., 2020). As a result, clustering may be driven by differences in mRNA stability or cytoplasmic transcript accumulation rather than current transcriptional state. For example, this may explain why only a single epithelial cluster was identified in the present snRNA-seq study, whereas at least two epithelial clusters were apparent in the scRNA-seq analysis. Likewise, fewer neuronal cell types could be confidently annotated in the present dataset compared to Krishnan et al. (2025). Technical factors may further contribute to discordance. Certain cell types, particularly large or fragile cells, may be lost during scRNA-seq tissue dissociation, which requires preservation of intact plasma membranes, whereas other cell types may be selectively lost or damaged during the freezing and nuclei isolation steps used in snRNA-seq protocols (Habib et al., 2017; Ding et al., 2020). Together, these biological and technical differences between nuclear and whole-cell RNA sequencing likely contribute to the incomplete concordance of cell-type clustering and marker gene expression observed between studies. In the absence of consistent in situ hybridization data many cluster annotations will, at present, remain elusive.

However, despite these limitations in precise cell-type annotation, several lines of evidence indicate that the clustering observed in this study reflects biologically meaningful structure rather than methodological artefact. First, multiple clusters exhibited distinct and internally consistent transcriptional signatures, including coherent expression of gene sets associated with major functional categories such as metabolism, structural components, and regulatory processes. Second, several clusters corresponded well to broadly defined anatomical or physiological cell types that were also identified in the previous scRNA-seq study, including ovary-associated cells, fat body and hemocyte-related populations, and epithelial and midgut-associated clusters, which also showed meaningful age-related changes in cell abundance and DE (see below). The reproducible detection of these major cell classes across independent studies using different methodologies supports the conclusion that the observed clustering captures genuine biological variation. Third, even in cases where direct correspondence between snRNA-seq and scRNA-seq clusters was incomplete, the presence of shared marker genes (even below significant enrichment) and related functional annotations suggests that these clusters represent related cellular states or subtypes rather than entirely unrelated or artifactual populations. Together, these observations indicate that, while fine-scale cell-type annotation remains challenging, the present snRNA-seq dataset provides a robust and biologically meaningful representation of cellular heterogeneity.

### Cluster abundance in young, old non-reproducing, and old reproducing Daphnia

Age- and reproduction-dependent differences in cluster composition indicate that aging in Daphnia involves structured remodeling of cellular states rather than uniform decline. Clusters putatively associated with ovary and fat body (clusters 0 and 5) are strongly enriched in young individuals and depleted in both classes of old animals, consistent with reduced reproductive and metabolic investment with age. In contrast, several clusters, including multiple with neural affinities and several unannotated populations, are relatively enriched in old individuals, suggesting either expansion of these cell populations or, more plausibly, age-associated shifts in transcriptional state that alter their representation in clustering space. Together, these patterns argue that aging is accompanied by a redistribution of transcriptional effort across tissues rather than a uniform attenuation of activity.

Reproductive status within old individuals modifies this redistribution in a selective manner. The enrichment of reproductively active old animals in a midgut-associated cluster (cluster 3), contrasted with the absence of such enrichment in a second midgut cluster (cluster 10), indicates that “rejuvenation” does not globally restore youthful cell composition. Instead, it appears to target specific functional compartments, particularly those associated with nutrient processing and metabolic support. This is consistent with the idea that restoration of reproductive function depends on selective reactivation of key physiological systems rather than wholesale reversal of aging.

Interpretation of cluster abundance changes is complicated by incomplete annotation and by the fact that clustering reflects transcriptional similarity rather than strictly defined cell types. Nevertheless, the magnitude, directionality, and reproducibility of these shifts across replicates strongly support their biological origin. In this light, the observed compositional changes are best understood as reflecting coordinated, system-level reallocation of cellular activity during aging and reproductive transitions, with only a subset of these changes being reversible.

### Cluster-specific differential expression

Differential expression within clusters reveals that aging in Daphnia is highly cell-type-specific and cannot be reduced to a single transcriptional trajectory. Instead, distinct clusters exhibit partially overlapping but clearly differentiated responses, involving structural components, metabolic pathways, and regulatory genes. For example, ovary-associated cluster 0 shows coordinated downregulation of structural proteins, mitochondrial components, and redox-related enzymes, alongside upregulation of transcription factors, cytochrome P450 enzymes, and transposon-derived transcripts. A related cluster (cluster 5) shares elements of this pattern but differs in key respects, including downregulation of lipid metabolism and germline-associated genes. These differences between closely related clusters indicate that aging does not act uniformly even within functionally related tissues, but instead reshapes transcriptional programs in a context-dependent manner.

Midgut-associated clusters exhibit a distinct transcriptional shift characterized by reduced expression of digestive enzymes and increased expression of membrane transporters and signaling components. This pattern is consistent with a transition away from active digestion toward altered nutrient handling or signaling states, although its functional consequences remain to be determined. In contrast, clusters with neural annotations show less consistent and more heterogeneous changes, suggesting either greater stability of transcriptional programs in neural tissues or unresolved heterogeneity within these clusters. Importantly, clusters lacking clear functional annotation nonetheless display coherent and recurrent aging-related changes, including downregulation of conserved signaling components and upregulation of cytochrome P450s and transposon-associated transcripts, supporting their biological relevance.

Across clusters, one of the most consistent features of aging is the increased expression of transposon- or retroelement-derived sequences, accompanied by changes in genes associated with mitochondrial function, oxidative stress, and transcriptional regulation. While the causal relationships among these processes remain unclear, their repeated co-occurrence across multiple cellular contexts suggests a general breakdown of transcriptional control coupled with activation of stress-responsive pathways. The absence of strong GO enrichment signals likely reflects both incomplete functional annotation in Daphnia and the distributed nature of these changes across pathways. Taken together, these results support a model in which aging proceeds as a mosaic of cell-type-specific transcriptional alterations, with limited convergence on a small set of shared pathways.

### Concordance with bulk RNAseq data: age-related DE

How do cluster-specific results reported here align with whole-body RNAseq comparison between young, old non-reproducing, and old reproducing *Daphnia* (Dua et al. 2025)? Generally there is a good concordance between bulk RNAseq and SN RNAseq results: a genes found to be up- or down-regulated in old, non-reproducing *Daphnia* in the bulk RNAseq experiment were more likely than by chance to be, respectively up-regulated in at least one cluster (one-tailed Fisher Exact Test P<0.0016) and down-regulated in at least one cluster (one-tailed Fisher Exact Test P<0.0001) in this study (Supplementary Data, tab ConcordanceWithBulk). Many of highlights of age-related DE reported in Dua et al. 2025 are represented in this category. These include 5 GMP synthases paralogs (up-regulated in old *Daphnia* in the bulk RNAseq data and in several clusters in this study, such as presumed midgut and epithelium) and several different genes of transposon origin (up-regulated in the bulk RNAseq data and up in several clusters in this study, including ovaries and fat body). It should be noted, that while some gene family members do show concordance between bulk and SN DE analyses, others, with similar expression rate, do not. It is not known whether this is caused by differences between experimental conditions or age at sampling, or between individual genes’ propensity to show DE expression in bulk vs. SN design, either due to differences in library preparation or in bioinformatics filtering and analysis.

On the other hand, several genes showed the opposite trends in the bulk- and SN- RNA seq data. In some cases this discrepancy could be readily explained by age-related changes in cell type abundance (Supplementary Table S6). One striking example of this discordance is the **ApoD** paralog 14.92 in contig 14F. In Dua et al. 92025) it was the only apolipoprotein D paralog expressed in young *Daphnia,* with ∼10-fold upregulation in the old samples and full reversal in reproductively “rejuvenated” ones. Here, the same paralog is slightly down-regulated in the ON samples in moderately abundant and ON-enriched cluster 2, indicating that the up-regulation detected in the bulk RNAseq data was due to cell type abundance change, not to cell-specific DE. The reverse pattern was observed in an **HSP-22** gene 0.4 in contig 292F, which was down-regulated in old bulk RNAseq samples, but is reported here to be up-regulated in cluster 5 (fat body), which is one of the clusters significantly reduced in abundance in the old samples (and down in two other clusters) and in several other genes (Supplementary Table S6).

### Concordance with bulk RNAseq data: reproductive “rejuvenation”-related DE

Similarly to the patterns described above, genes with reported DE between old, non-reproducing and old, reproducing *Daphnia* were significantly more likely than by chance to be discovered as, being concordantly, up- or down- regulated in one or more cell clusters in this study (FET P<0.016 and P<0.0001, respectively, for up- and down-regulated genes). The unidentified cluster 11 stood out with this respect, showing concordant down-regulation of several potentially rejuvenation-related genes, including genes encoding two different serine protease, a cadhedrin ortholog, a neurotropin ortholog, a dusky-like protein (an ortholog of *Drosophila **dusky*** known to regulate epidermal cell shape), a vitamin D 25-hydroxylase/ cyt P450, a heme-binding protein, a chitin deacetylase, a Zn-binding domain containing protein, a Zn-finger protein, a neprilysin ortholog, and several proteins with unknown functionality or orthology. Several other clusters showing DE in old, reproducing *Daphnia* relative to old, non-reproducing ones (clusters 1, 2, 6, and 7) also escaped an unambiguous identification. An in-situ hybridization study is necessary to establish the location and nature of this cell type with the potential of revealing cellular basis of reproductive rejuvenation. One cluster with positive identification, however, was also showing a concordant DE with the bulk RNAseq data, and that was, not surprisingly, cluster 0 identified as ovaries. It showed an up-regulation of a homeobox protein homologous to *Drosophila **apterous***, a small ubiquitin-related modifier protein, and a tRNA-uracil-methyltransferase, all of which showed a concordant up-regulation during “rejuvenation”.

## Conclusions

We were able to identify several cell clusters that show either change in abundance, or DE, or both, in a comparison of old *Daphnia* to young *Daphnia*, and in a comparison of old, reproductively “rejuvenated” *Daphnia* to old, reproductively senescent ones. Some of these changes amounted to the reversal (or prevention) of age-related changes, often with a significant concordance with previously published bulk RNAseq data. Cell clusters with the strongest age-related and “rejuvenation”-related transcript- or cell abundance changes include ovaries, midgut, and several clusters with yet unknown functionality.

## Data Accessibility

Raw reads are available at NCBI SRA, Accession numbers SRR32324292 - SRR32324299; processed files at NCBI GEO, Accession number GSE317254.

## Acknowledgements

We are grateful to Dieter Ebert for providing reference genome and transcriptome ahead of public release, to Leon Peshkin and Marc Kirschner for valuable advise and discussion and to Ishan Dua, Ashit Dutta, and Meridith Smith for laboratory assistance.

## Supplementary materials

## Supplementary Tables

**Supplementary Table S1.** Results of doublet-removal procedure (scDblFinder; min_cells = 3; dbr_used = 0.05).

| sample_id | n_cells | n_singlets | n_doublets |
| --- | --- | --- | --- |
| Yo2 | 2063 | 1912 | 151 |
| Yo3 | 2016 | 1863 | 153 |
| Yo4 | 1155 | 1048 | 107 |
| Yo5 | 3456 | 3130 | 326 |
| On1 | 1060 | 966 | 94 |
| On2 | 1378 | 1229 | 149 |
| Or1 | 809 | 730 | 79 |
| Or2 | 1938 | 1754 | 184 |

**Table S2.** Results of cycling over the parameters of the number of PCA dimension to be used (X) and resolution (res).

| dims=1..X | res | Number of clusters | Number of cluster into which the 0+5 supercluster is split |
| --- | --- | --- | --- |
| X=20 | 0.4 | 16 | 2 |
|  | 0.6 | 18 | 3 |
|  | 0.8 | 20 | 3 |
|  | 1 | 21 | 3 |
|  | 1.2 | 26 | 5 |
| X=25 | 0.4 | 17 | 2 |
|  | 0.6 | 20 | 3 |
|  | 0.8 | 21 | 3 |
|  | 1 | 22 | 4 |
|  | 1.2 | 24 | 4 |
| X=30 | 0.4 | 17 | 2 |
|  | 0.6 | 19 | 3 |
|  | 0.8 | 21 | 3 |
|  | 1 | 23 | 4 |
|  | 1.2 | 23 | 4 |
| X=40 | 0.4 | 17 | 2 |
|  | 0.6 | 20 | 3 |
|  | 0.8 | 22 | 3 |
|  | 1 | 24 | 4 |
|  | 1.2 | 27 | 4 |
| X=50 | 0.4 | 16 | 1 |
|  | 0.6 | 18 | 2 |
|  | 0.8 | 23 | 3 |
|  | 1 | 25 | 4 |
|  | 1.2 | 27 | 5 |

**Supplementary Table S3.** Contingency table analysis of cluster composition. Clusters column includes cluster IDs and preliminary annotations, unless ambiguous. Full color: significant in an sample-level negative binomial generalized linear model (NB-GLM test, FDR<0.1. Diluted color: significant in a contingency test FDR<0.001 without accounting for batch effects.

| Count<br>Expected<br>Cell Chi^2 | Samples: |  |  | Total | Enrichment |
| --- | --- | --- | --- | --- | --- |
|  | On | Or | Yo |  |  |
| Cluster 0<br>Ovary | 218<br>495.0927<br>155.0828 | 669<br>560.3375<br>21.0722 | 1963<br>1794.57<br>15.8081 | 2850 | Yo and Or vs. On |
| 1 | 424<br>378.8762<br>5.3742 | 326<br>428.8056<br>24.6475 | 1431<br>1373.318<br>2.4227 | 2181 | On vs. Or |
| 2 | 380<br>254.1476<br>62.3214 | 270<br>287.6399<br>1.0818 | 813<br>921.2125<br>12.7115 | 1463 | On vs. Yo |
| 3<br>Midgut | 146<br>224.442<br>27.4153 | 360<br>254.0196<br>44.2164 | 786<br>813.5383<br>0.9322 | 1292 | Or vs. On |
| 4<br>Epithelium | 213<br>210.371<br>0.0329 | 287<br>238.0943<br>10.0455 | 711<br>762.5348<br>3.4829 | 1211 | Or vs. Yo |
| 5<br>Fat body | 67<br>181.3603<br>72.1121 | 26<br>205.2605<br>156.5538 | 951<br>657.3793<br>131.1467 | 1044 | Yo vs. both O |
| 6 | 120<br>112.3947<br>0.5146 | 107<br>127.2064<br>3.2097 | 420<br>407.3988<br>0.3898 | 647 | None |
| 7 | 150<br>96.23907<br>30.0319 | 111<br>108.9217<br>0.0397 | 293<br>348.8392<br>8.9383 | 554 | On vs. Yo |
| 8<br>Neural | 158<br>68.79183<br>115.6838 | 75<br>77.85741<br>0.1049 | 163<br>249.3508<br>29.9035 | 396 | On vs. Yo |
| 9<br>Possibly<br>neural | 59<br>51.42015<br>1.1173 | 25<br>58.19645<br>18.9359 | 212<br>186.3834<br>3.5208 | 296 | Yo vs. Or |
| 10<br>Midgut | 55<br>30.92158<br>18.7497 | 56<br>34.99651<br>12.6054 | 67<br>112.0819<br>18.133 | 178 | Or and On vs. Yo |
| 11 | 96<br>20.67229<br>274.4865 | 23<br>23.39655<br>0.0067 | 0<br>74.93116<br>74.9312 | 119 | Or vs. Yo |
| 12<br>Neural | 30<br>19.62999<br>5.4782 | 26<br>22.21689<br>0.6442 | 57<br>71.15312<br>2.8152 | 113 | On vs. Yo |
|  | 24 | 68 | 19 |  | Or vs. Yo |
|  | 19.28256 | 21.82367 | 69.89377 |  |  |
| 13 | 1.1541 | 97.7037 | 37.0588 | 111 |  |
|  | 12 | 12 | 50 |  | None |
| 14 | 12.85504 | 14.54911 | 46.59585 |  |  |
| Follicle | 0.0569 | 0.4466 | 0.2487 | 74 |  |
|  | 22 | 37 | 10 |  | On and Or vs. Yo |
| 15 | 11.98645 | 13.56606 | 43.44748 |  |  |
| Neural | 8.3654 | 40.4796 | 25.7491 | 69 |  |
|  | 19 | 4 | 3 |  | On and Or vs. Yo |
|  | 19.2826 | 21.8237 | 69.8938 |  |  |
| 16 | 1.1541 | 97.7037 | 37.0588 | 26 |  |
| Total | 2193 | 2482 | 7949 | 12624 | Total |

**Supplementary Table S4.**
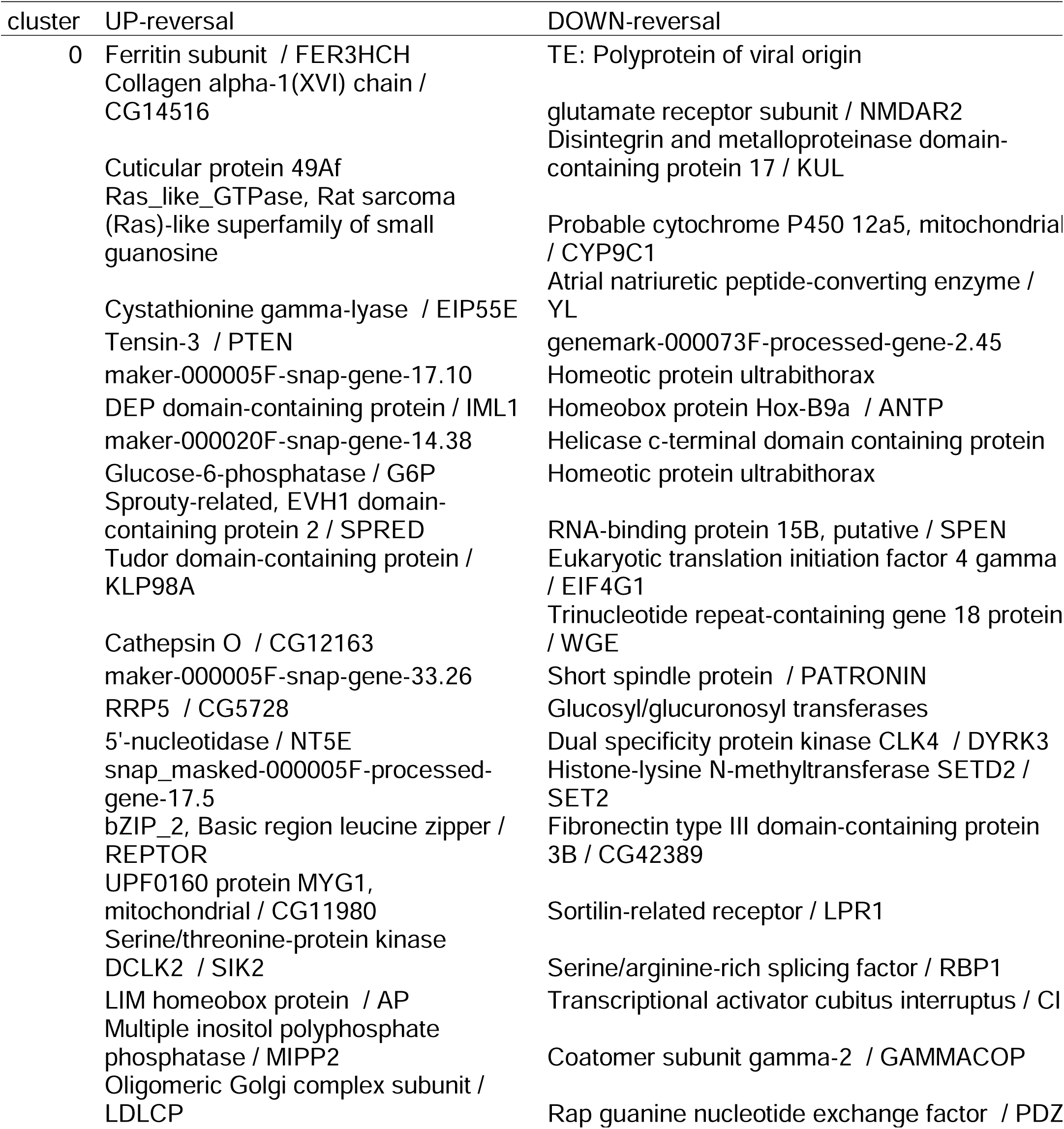
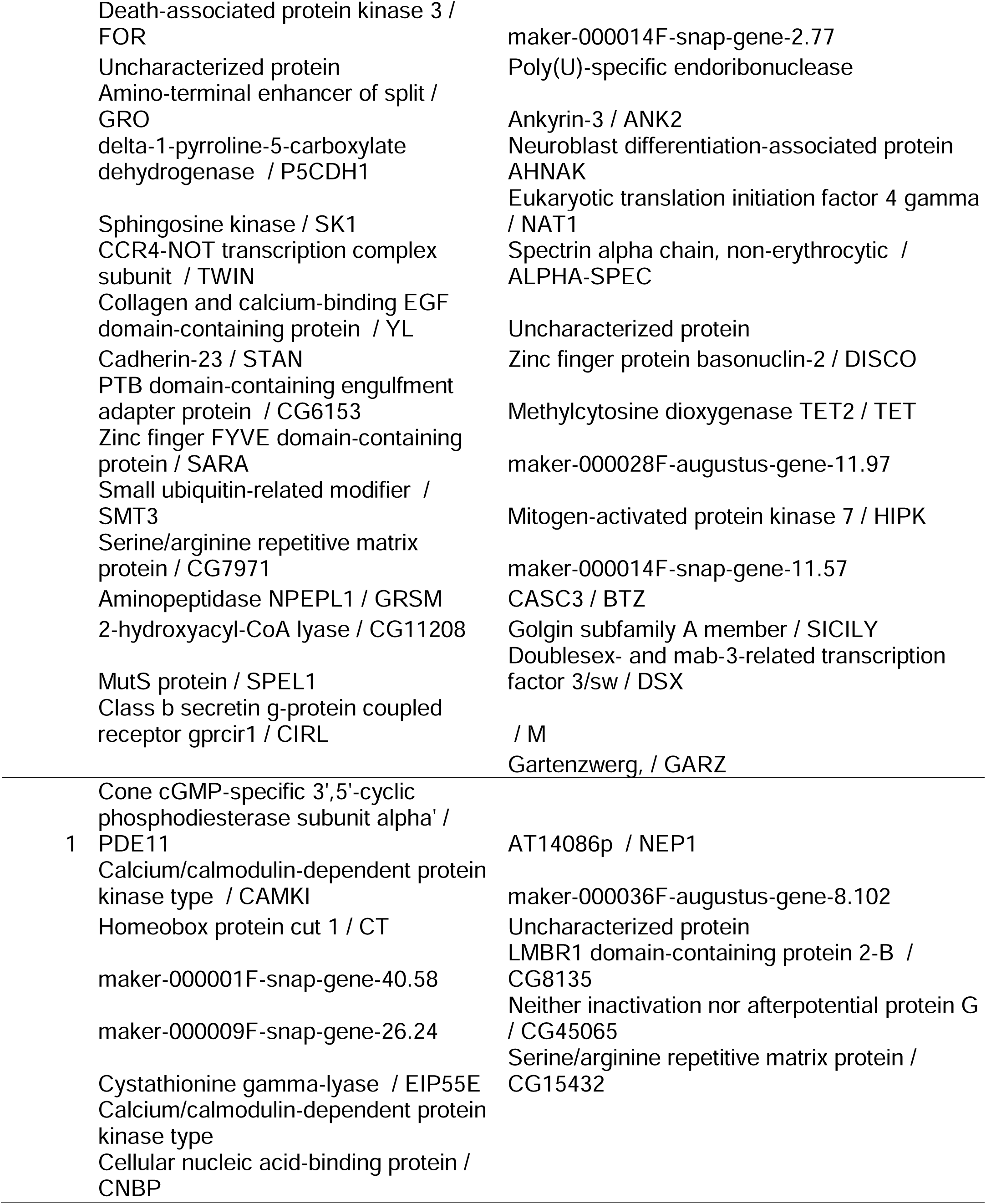
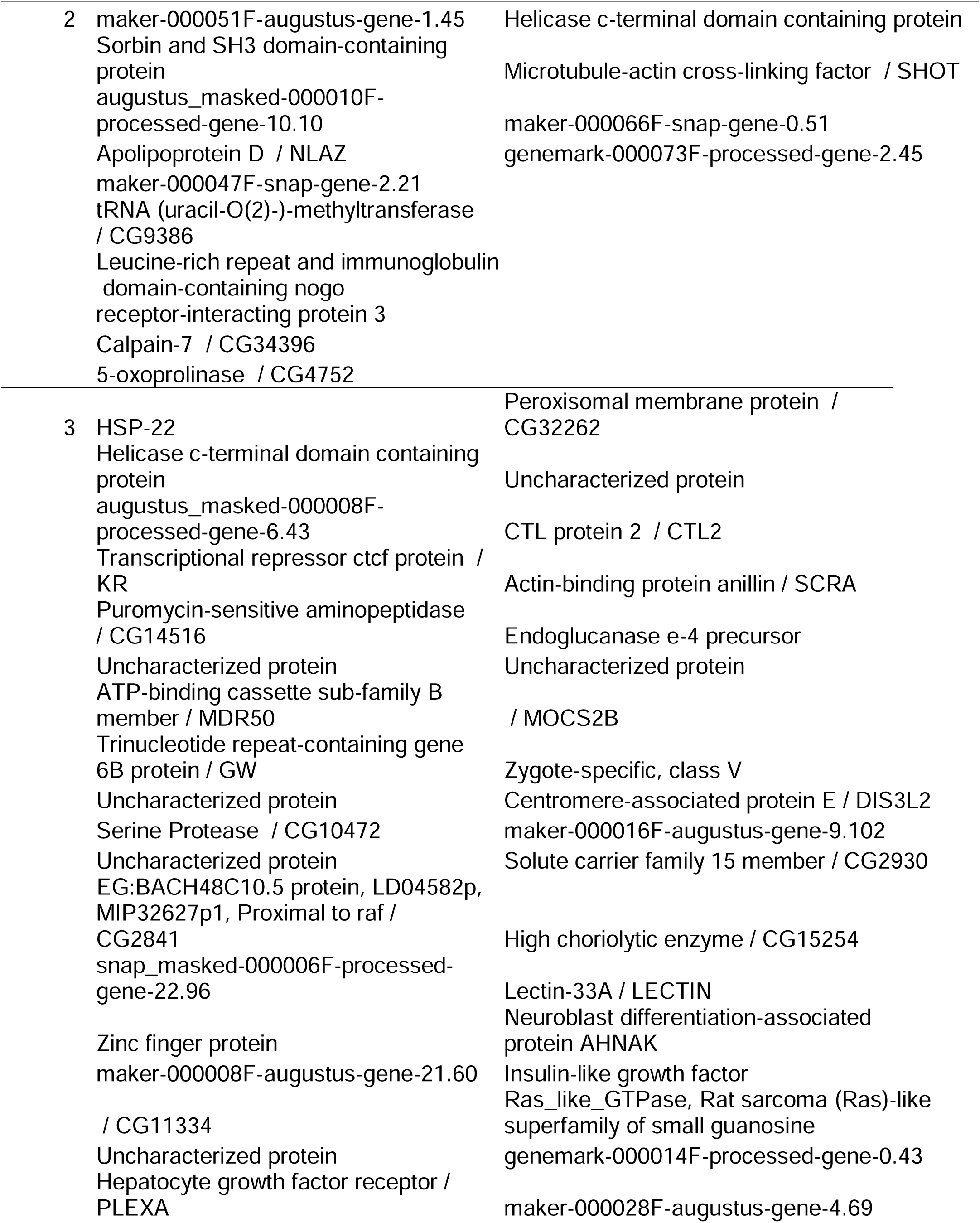
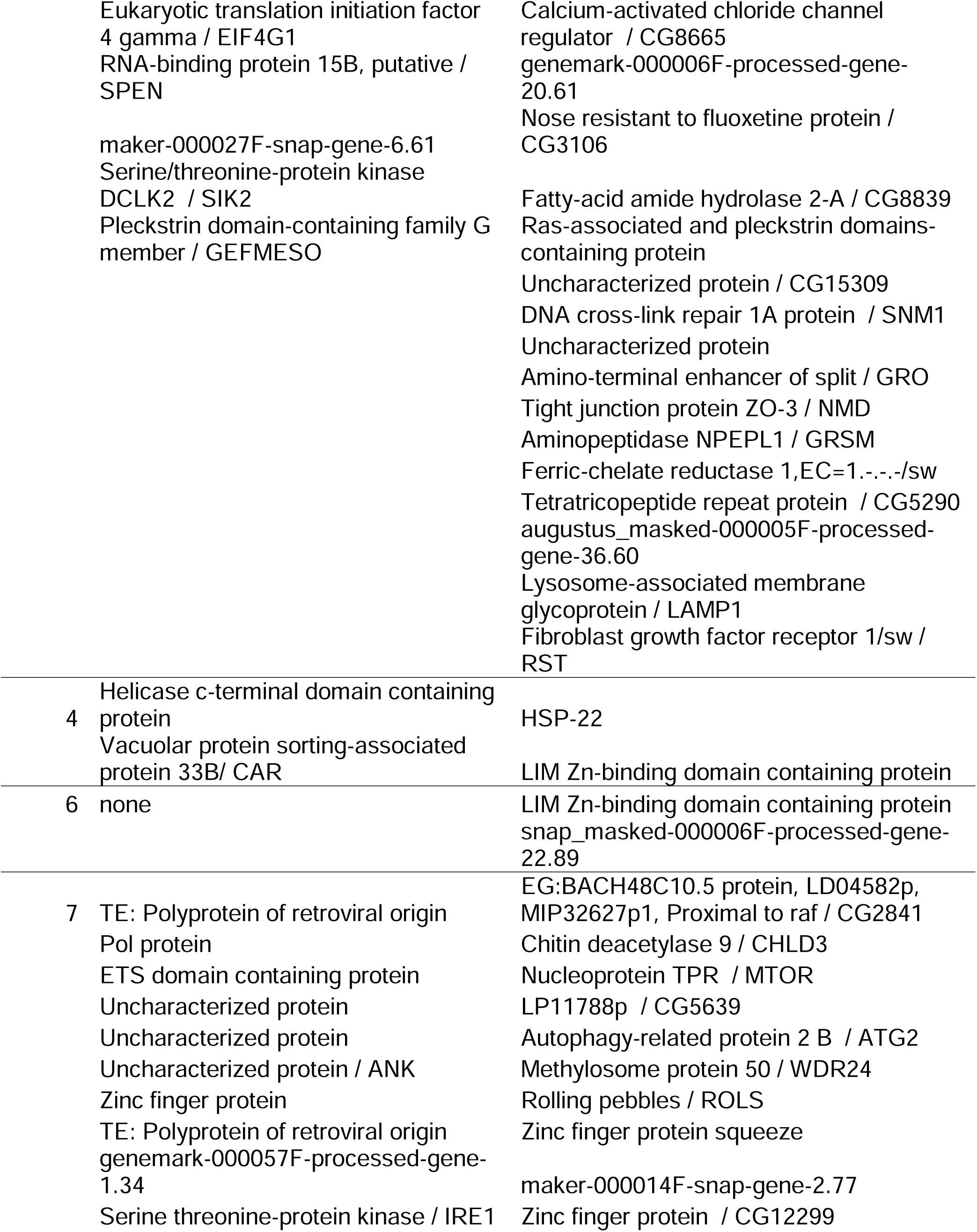
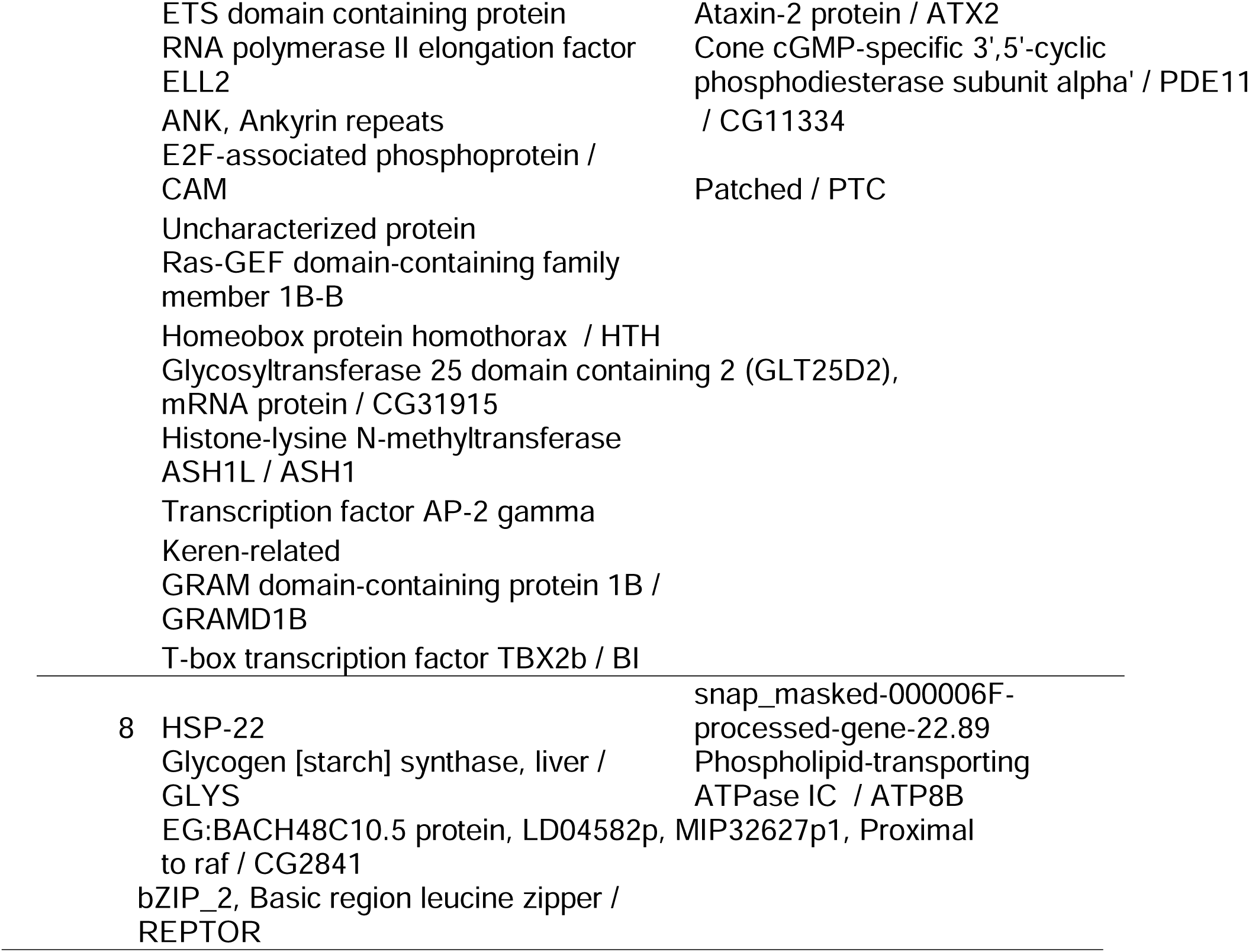
Genes showing reversal of age-related changes in reproductively “rejuvenated” *Daphnia. “*UPreversal”: down-regulated in old, non-reproducing *Daphnia* relative to young ones, but up in old, reproducing *Daphnia* relative to old, non-reproducing ones; “DOWN-reversal”: Up in old, non-reproducing *Daphnia* relative to young ones, but down-regulated in old, reproducing *Daphnia* relative to old, non-reproducing ones.

**Supplementary Table S5.** Genes showing co-directional age- and “rejuvenation”-related changes.

| cluster | both Up | both Down |
| --- | --- | --- |
| 0 | none | Maelstrom / MAEL<br>Mesoderm induction early response protein |
| 1 | Cell adhesion molecule / DIP-beta<br>Steroid 17-alpha-hydroxylase/17,20 lyase / CYP18A1<br>maker-000002F-snap-gene-7.0<br>Two pore calcium channel protein 2 | Major Facilitator Superfamily Transporter / MFS3<br>Vacuolar protein sorting-associated protein 33B/ CAR |
| 2 | phosphatase and actin regulator<br>Zinc finger C2HC domain-containing protein 1A<br>StAR-related lipid transfer 13 protein | Transcription factor capicua / CIC<br>Contactin associated protein like 5-1 / AXO<br>Solute carrier family 22 member<br>Vacuolar protein sorting-associated protein 33B / CAR |
| 3 | none | PIF1-like helicase |
| 4 | genemark-000005F-processed-gene-33.77 | Zinc finger and SCAN domain-containing protein / CROL<br>Cuticle N-domain containing protein<br>maker-000051F-augustus-gene-1.45 |
| 6 | none | maker-000051F-augustus-gene-1.45 |
| 7 | none | Steroidogenic Acute Regulatory protein<br>polyprotein of retroviral origin<br>TE: Polyprotein of retroviral origin<br>maker-000007F-snap-gene-12.43 |

**Supplementary Figure S1.**
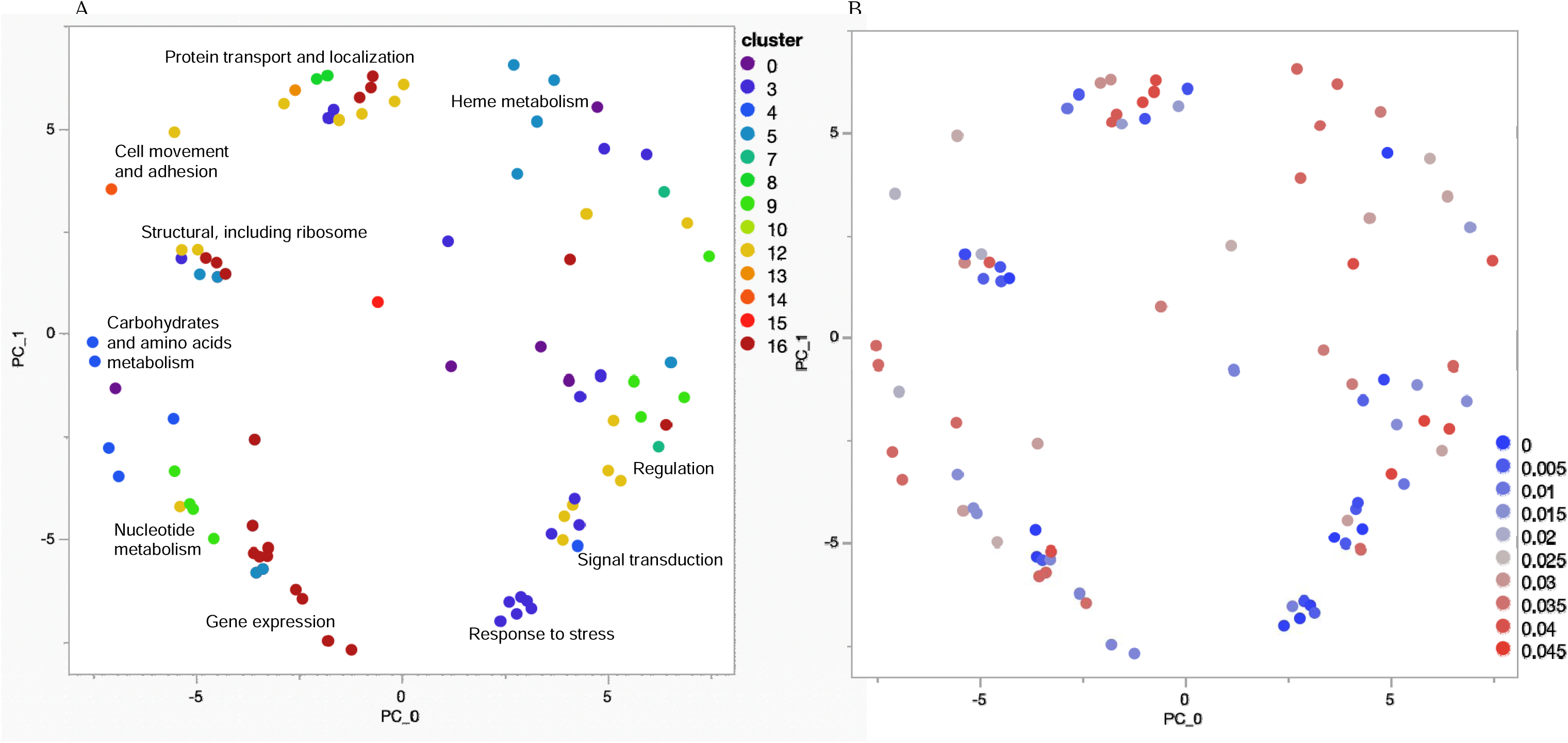
ReViGO plots showing semantics of cluster composition. A: semantic terms colored by cluster color. B: semantic terms colored by association strength.

**Supplementary Fig. S2.**
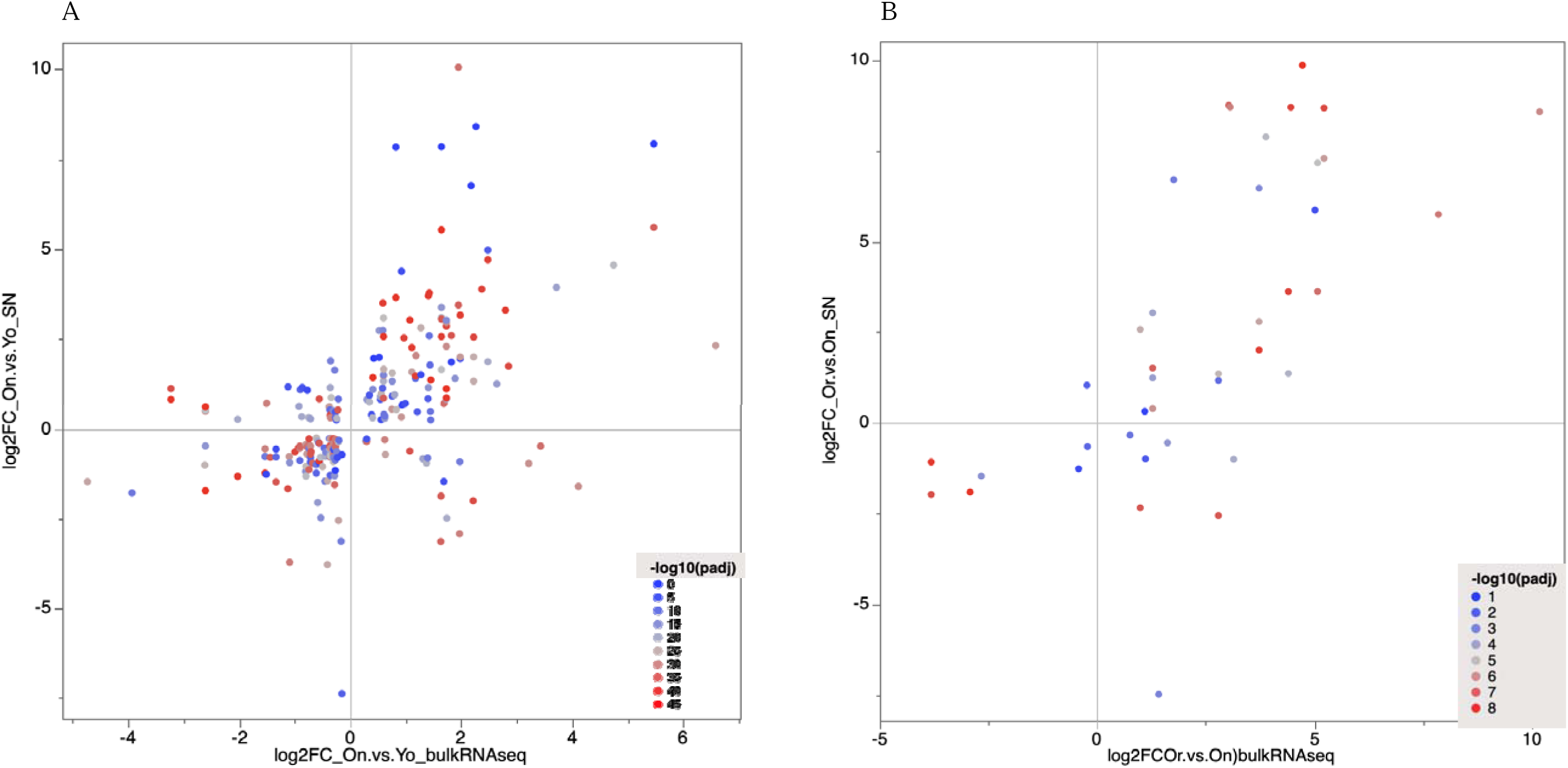
Concordance between DE estimates in bulk RNAseq study (Dua et al. 2025; log2(FC), horizontal axis) and this study (DE in at least 1 cluster; log_2_(FC), vertical axis). A: comparison of old, non-reproducing *Daphnia* to young *Daphnia.* B: comparison of old, reproducing *Daphnia* to old, non-reproducing *Daphnia.* Each dot is a gene-cluster combination in this study; colored by -log_10_(p_adj) for a given cluster. Only genes with p_adj<0.1 are shown.

## Notes

https://www.ncbi.nlm.nih.gov/bioproject/PRJNA1222346

### Competing Interest Statement

The authors have declared no competing interest.

### Summary of Updates

Improved Introduction and Discussion, corrected several minor issues.

### Funding

- Impetus Foundation

